# Global genome decompaction leads to stochastic activation of gene expression as a first step toward fate commitment in human hematopoietic stem cells

**DOI:** 10.1101/2020.09.09.289751

**Authors:** Parmentier Romuald, Moussy Alice, Chantalat Sophie, Racine Laëtitia, Sudharshan Ravi, Papili Gao Nan, Stockholm Daniel, Corre Guillaume, Fourel Geneviève, Deleuze Jean-François, Gunawan Rudiyanto, Paldi Andras

## Abstract

When human cord blood derived CD34+ cells are induced to differentiate *in vitro*, they undergo rapid and dynamic morphological and molecular transformations that are critical for fate commitment. Using ATAC-seq and single-cell RNA sequencing, we detected two phases in this process. In the first phase, we observed a rapid and global chromatin opening that makes most of the gene promoters in the genome accessible, followed by widespread upregulation of gene transcription and a concomitant increase in the cell-to-cell variability of gene expression. The second phase is marked by a slow chromatin closure and a subsequent overall downregulation of gene transcription and emergence of coherent expression profiles corresponding to distinct cell subpopulations. These observations are consistent with a model based on the spontaneous probabilistic organization of the cellular process of fate commitment.

## Introduction

Fate commitment of hematopoietic cells has been widely studied and is commonly considered as a paradigm of cell differentiation in general. Traditionally, differentiation is believed to proceed through a series of binary fate decisions under the action of key instructive factors inducing specific changes in the cell that lead to stepwise switches of the expression profiles at critical decision points (Kawamoto and Katsura, 2009). The typical representation of this process is a hierarchical decision tree. Such a strict hierarchical process must imply tight regulation of gene expression. A number of genes that play a key role in the process have been identified (Sive and Göttgens, 2014). But recent observations challenge the assumption of a strictly ordered process. Single-cell gene expression studies demonstrated that, soon after their stimulation for differentiation, multipotent CD34+ cells go through a phase of disordered gene expression called “multilineage primed” phase characterized by concomitant expression of genes typical for alternative lineages (Hu et al., 1997; Moussy et al., 2017; Nimmo et al., 2015; Pina et al., 2012). Other studies demonstrated that hematopoietic stem cells (HSC) gradually acquire lineage characteristics along multiple directions without passing through discrete hierarchically organized and demarcated progenitor populations (Velten et al., 2017). Instead, unilineage-restricted cells emerge directly from a continuum of low-primed undifferentiated hematopoietic stem and progenitor cells (Velten et al., 2017). This phase is accompanied by instabilities and fluctuation of the cell transcriptome, morphology and dynamic cell behavior (Moussy et al., 2019, 2017). How this quasi-random gene expression pattern is generated and how it transforms into a defined gene expression profile remain unknown. In order to answer these questions, we investigated the nature, the order and the timescale of the early chromatin and transcriptional changes that follow the induction of differentiation in CD34+ cells.

To do this, we performed single cell RNA sequencing of human cord blood CD34+ cells at different time points during the 96h period following their stimulation, a period that has been shown to be critical for cell fate decision (Moussy et al., 2017). The gene expression profiles were correlated to the DNA accessibility changes determined by ATAC-seq at defined time-points during the same period. The data revealed strikingly different dynamics for chromatin accessibility and gene expression that challenges the classical model based on specific stepwise switches.

## Results

### Initial transcription burst precedes stable expression profiles

The experimental strategy is shown in **Figure 1A**. Human CD34+ cells were isolated from the cord blood of two healthy donors and cultured in the presence of early acting cytokines as described previously (Moussy et al., 2017). To identify the transcriptional signatures and to estimate their variability at the earliest stages of the differentiation process, we performed MARS-seq (massively parallel single-cell RNA-sequencing, see Materials and Methods) on CD34+ cells randomly sorted at different time points (5h, 24h, 48h, 72h and 96h) after the cells were cultured in the presence of cytokines (Jaitin et al., 2014). The uniform random sampling of a heterogenous population allowed us to evaluate the global changes without any preconceived ideas on the cell categories present in the population. The quantification of gene expression was calibrated using unique molecular identifier (UMI) marked RNAs. Details about quality control of the results are shown in **Table S1.** In order to avoid potential bias due to batch correction, the results of the two donors were analyzed separately.

**Figure 1.**
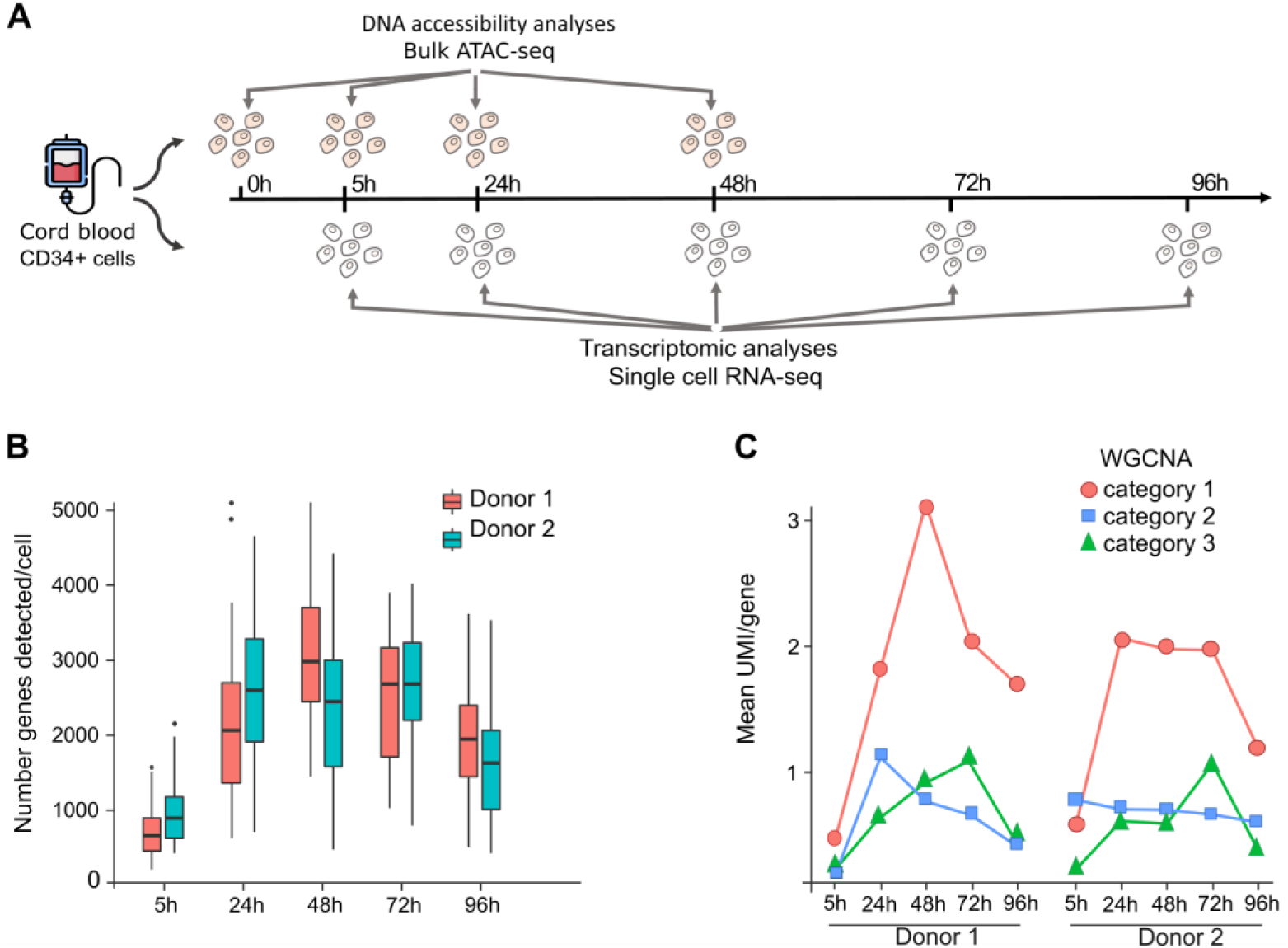
Gene expression dynamics of cord blood derived CD34+ cells. (A) CD34+ cells were isolated from human cord blood and cultured in serum-free medium with early acting cytokines. Single-cell RNA sequencing (scRNA-seq) was used to analyze single-cell transcription at 5h, 24h, 48h, 72h and 96h. Concomitantly, at 0h, 5h, 24h and 48h, 5000 living cells were collected to perform ATAC-seq protocol in order to study DNA accessibility dynamics. (B) Number of detected genes per cell with scRNA-seq. Two donors were analyzed separately, both showed similar dynamics. For the Donor 1 see the Results section. For Donor 2, at 5h - 760 genes +/- 297, at 24h – 2298 genes +/- 822, at 48h – 2036 genes +/- 809, at 72h – 2217 genes +/- 612 and at 96h – 1420 genes +/- 630. (C) Weighted correlation network analysis (WGCNA) reveals groups of genes with similar dynamic patterns in the average mRNA expression in Donor1 and Donor2. Note that group 1 reproduces the dynamic pattern observed for genes showing detectable expression in single cell in (B). Category 1 = 5194 genes (Donor1) and 5518 genes (Donor2), group 2 = 3977 genes (Donor1) and 2602 (Donor2), group 3 = 1089 genes (Donor1) and 609 genes (Donor2).

The results revealed important features in global gene expression dynamics (**Figure 1**). Following stimulation, the transcriptome underwent rapid and substantial quantitative and qualitative changes. Both the number of expressed genes per cell and the number of mRNA molecules per gene increased substantially (**Figure 1B and 1C**). The average number of genes detected per cell at 5h was only 512+/-243 in Donor1. This number increased to 1693 +/-813 at 24h and 2543+/- 751 at 48h, but then decreased to 2014+/-714 at 72h and to 1612 +/-613 at 96h. Numbers for Donor 2 were very similar (see legend **Figure 1B**). The rapid increase in global transcription activity occurred mainly during the first 48h, suggesting that cells expand their repertoire of transcribed genes during the initial phase of the differentiation process.

Examination of individual genes confirmed a corresponding increase in the number of mRNA molecules. Categories of genes with highly correlated mean expression patterns over time could be defined using Weighted Correlation Network Analysis (WGCNA) (**Figure 1C**), and the three largest categories together sum up to more than 10200 genes for Donor 1 and 8700 genes for Donor 2. Strikingly, all three categories display a similar time profile with an initial increase followed by a subsequent decrease, pointing to a genome-wide phenomenon. Thus, an average CD34+ cell responds to cytokine stimulation with a strong, but transient gene upregulation, both in terms of gene number and number of transcripts. During the 24h to 48h period after stimulation, the gene fraction transcribed in each individual cell rose to reach approximately 10-15% of all genes in the genome (**Figure 1B**). After 72h, this number started to decrease, coinciding with the time when the first signs of lineage-specific transcriptional changes appear (Moussy et al., 2017).

In order to detect emerging gene expression patterns and characterize the lineage progression and the possible trajectories of the cells during the period under scrutiny, we analyzed our single-cell RNA dataset using CALISTA (Clustering And Lineage Inference in Single-Cell Transcriptional Analysis) [10]. CALISTA is likelihood-based method that uses the two-state stochastic model of gene transcription to describe the cell-to-cell variability of gene expression at single-cell level (Peccoud and Ycart, 1995). CALISTA can be used to identify cell clusters and cell lineages, calculate single-cell transcriptional uncertainty and assign to each cell a likelihood value which reflects the joint probability of its gene expression levels (mRNA counts). Since we were interested in general tendencies in transcription changes, we analyzed the single-cell mRNA datasets from two donors independently. In this way, biases related to batch effects and their corrections can be avoided. For both donors, CALISTA identified five single-cell clusters on the basis of the 200 most variable genes (**Table S2**). In both donors, clusters #1 and #2 were essentially composed of cells isolated at 5h and 24h, respectively (**Figure 2B and S1B**). Clusters #3, #4 and #5 were mixed containing cells collected at 48h, 72h and 96h (**Figure 2B and S1B**). This suggests that individual cells progress at their own pace. Some cells reached the profile corresponding to clusters #4 or #5 as early as 48h, while others needed 96h to do so.

**Figure 2.**
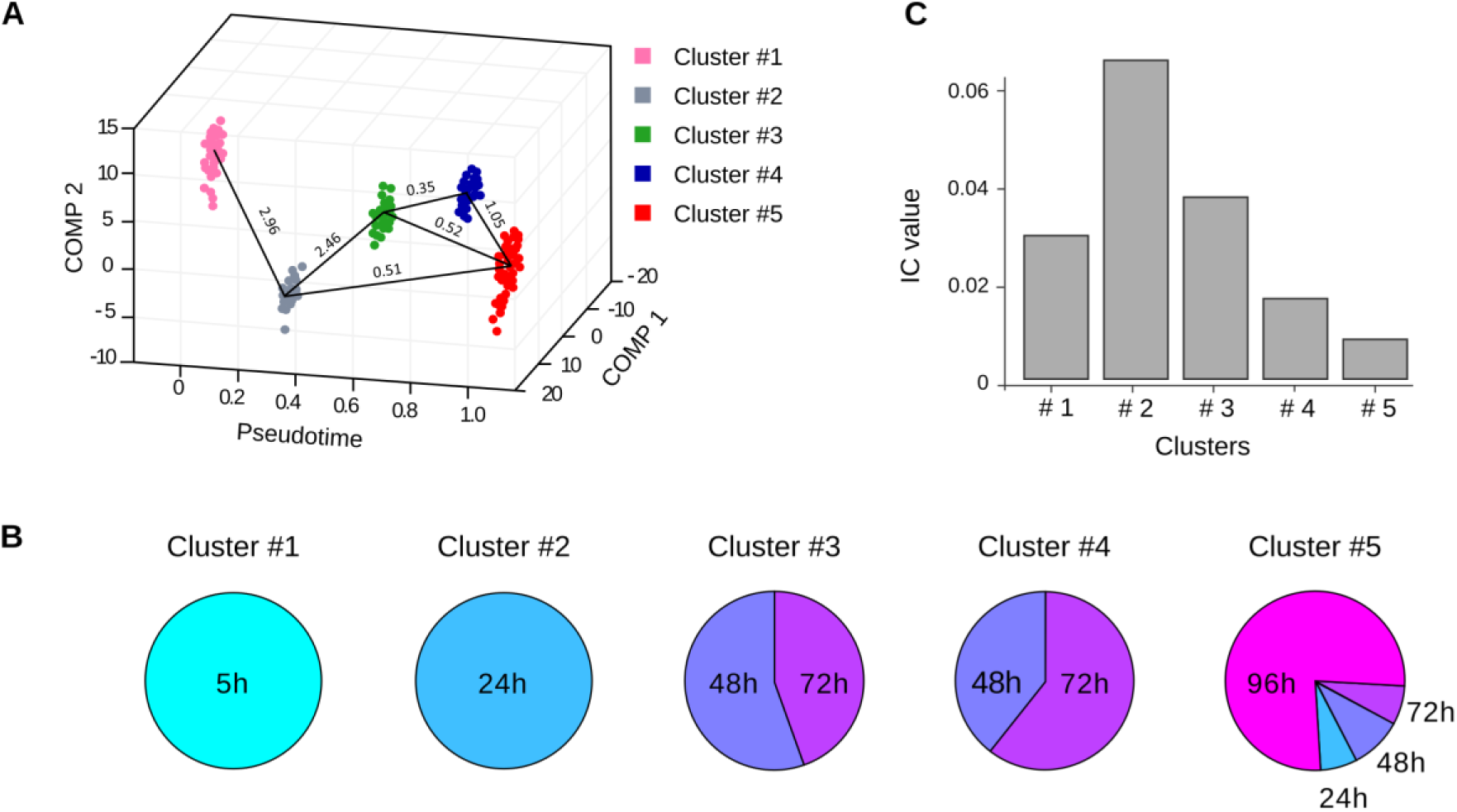
Evolution of transcriptome profiles after cell stimulation in Donor 2. (A) Transcriptome clusters identified by CALISTA for Donor 2. Each dot corresponds to a cell in the single-cell transcriptomic dataset sampled at 5h, 24h, 48h, 72h and 96h. The x axis corresponds to the pseudotime, the y-z axes to the first and second principal component (PC). The color code is given in the upper right inset. The transition edges are represented by black plain lines between the clusters and the numbers are “cluster distances”, a likelihood-based measure of dissimilarity (distance) between cell clusters. Note that there are several ways a cell can reach the clusters 3 to 5. The results for Donor 1 are shown on **Figure S1**. (B) Contribution of the cells collected at different time points to the clusters identified by CALISTA. The mixed composition of the clusters #3 to #5 may reflect the different rates of cell transformation and the multiplicity of cell trajectories. (C) I_c_ index calculated for each cluster of Donor 2 as described in (Mojtahedi et al., 2016). The maximum is reached for cluster #2, indicating a phase of critical transition at 24h. After 24h, cells from cluster #3, #4 and #5 undergo stabilization, leading to a decreasing I_c_index value.

CALISTA also produces “cluster distances” between each pair of clusters based on the maximum difference in the cumulative likelihood values of the gene expression distribution (Papili Gao et al., 2020). This index helps to visualize the most likely sequence of the lineage progression (**Figure 2A** and **S1A**). Overall, the two graphs show highly similar lineage trajectories. Importantly, the close distances between the clusters #3, #4 and #5 makes likely that a cell can reach any of these clusters through different pathways or switch between them, as suggested by the reported time-lapse observations (Moussy et al., 2017).

In order to detect early-warning signals that would indicate cell state transitions, we calculated for each cluster the “index for critical transitions” (I_c_) as described in (Mojtahedi et al., 2016). To do this, we calculated the pairwise gene-gene correlation between all pairs of gene vectors (R(g_n_,g_m_)) and the cell-cell correlation between all pairs of cell state vectors (R(c_i_,c_j_)). The analysis was performed separately for each cluster and each donor. Only the correlations with a Pearson coefficient higher than 0.70 were taken into account. The I_c_ is calculated as the ratio between the average of all R(g_n_,g_m_)-s and R(c_i_,c_j_)-s (Mojtahedi et al., 2016). The results shown on **Figure 2C and S1C** indicate that in both donors the I_c_ sharply increased towards a maximum between 24h and 48h and decreased by 72h to 96h (**Figure 2C and S1C**) - a typical hallmark of a critical transition state.

Then, we performed a comparative Gene Ontology analysis of the cell clusters. For this, we used the list of genes for which the pairwise gene-gene correlation score was greater than 0.70. The top “molecular function” GO categories (p < 0,01) were compared between the clusters (**Figure S2**) using the “compareCluster” function of the Cluster Profiler package (Yu et al., 2012). The analysis showed similar enriched GO terms among clusters for donor 1 and donor 2. Cluster #1 is characterized essentially with broad-spectrum terms associated to translation, transcription activities and cellular interaction. These categories constitute a common base for all clusters. Cluster #2 and #3 showed the greatest variety of enriched GO terms, ranging from nucleotide synthesis to metabolic activities, but with no apparent cell type related functions. Finally, in cluster #5, GO terms pointing to erythroid lineage related functions emerged (see **Table S3** for GO terms enrichment statistics), suggesting that these cells are progressing in their lineage commitment.

In order to reveal potentially active regulatory interactions that could account for the transcription dynamics, we explored on a global scale the correlation between changes in the expression of transcription factor-coding genes (TFs) and changes in the expression of their target genes. To do so, we sorted the genes according to the evolution of their mRNA levels. This classification is based on the number of UMIs detected in a cell (see STAR Materials and Methods section for details). Genes that showed a statistically significant change in the corresponding mRNA level in the two donors are referred to as differentially expressed (DE). Focusing on the early changes, out of the total number of 14,045 genes that were expressed in at least one time point, we found 5,274 DE genes between 5h and 24h. Note that such DE genes were mainly upregulated, as only 110 genes were found downregulated. Genes with unchanged or undetected mRNA level were designated as “non-DE”. We found 8,771 non-DE genes between 5h and 24h. Among the 470 expressed genes encoding transcription factors (TFs), 56 showed a significant change in expression between 5h to 24h., labeled as DE-TF genes. Gene targets of the DE-TFs were identified using the Regulatory Circuits resource (Marbach et al., 2016). We found 4415 potential DE target genes for the 470 TFs. Finally, among them, the target genes of the 56 DE-TFs are overrepresented (enriched). Indeed, 2,630 of the target DE-genes (60% of 4415) are targeted by at least one of the DE-TFs (p=1.4×10^−6^) p=1.4×10^−6^, two-sided Fisher exact test) (**Figure 3A**).

**Figure 3.**
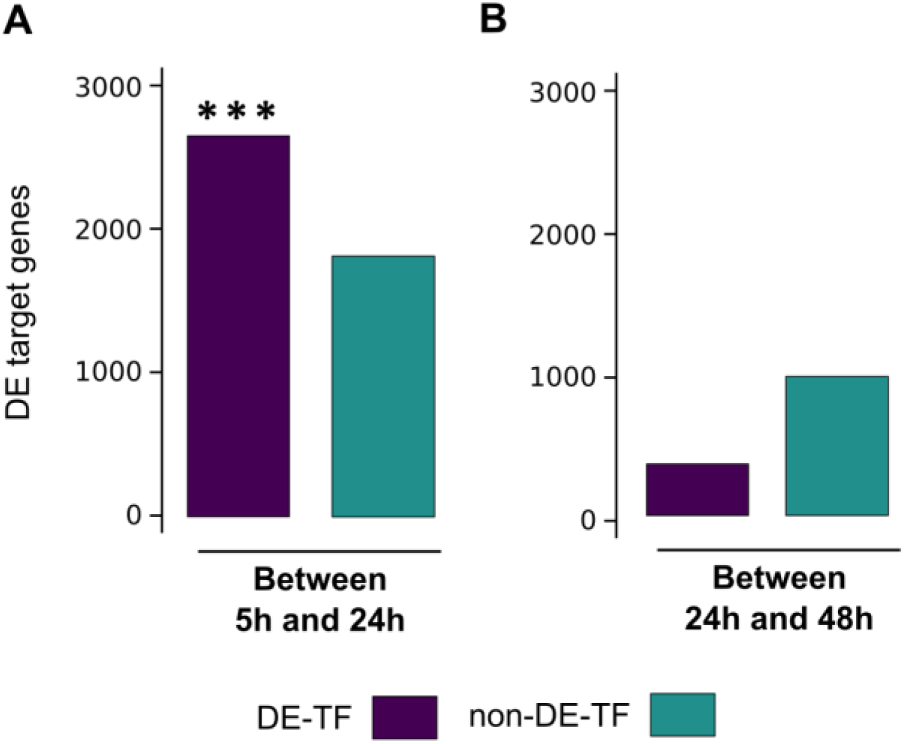
Global influence of transcription factors on targeted gene expression. (A) Total number of differentially expressed target genes (DE target genes) whether their associated TF is differentially expressed (DE-TF), or non-differentially expressed (non-DE-TF) between 5h and 24h. (B) Total number of differentially expressed target genes (DE target genes) whether their associated TF is differentially expressed (DE-TF), or non-differentially expressed (non-DE-TF) between 24h and 48h. Note that the association between differential expression (DE target genes) and the differentially expressed TF-coding genes targeting them is significant only between 5h and 24h.

These observations suggest that the increase of transcription between 5h and 24h could be potentially facilitated by the activity of TFs targeting them. However, TF activity is not sufficient to fully explain the expression changes since 40% of the DE target genes increase their transcription without being targeted by a DE-TF. The same strategy applied to the period between 24h to 48h revealed a different dynamic compared to the first-time interval (5h to 24h). First, only 16 TFs were detected as DE and among them, 8 were already classified as such between 5h and 24h. This decrease is expectedly accompanied with a drastic drop of the number of DE target genes from 4415 (5h-24h) to 1259 (24h-48h).

### Chromatin decompaction is a non-specific response to cell stimulation

In order to uncover if global chromatin changes occur during the critical state transition period, we determined the DNA accessibility in the CD34+ cells of three independent donors using ATAC-seq (Corces et al., 2016) at four time points (0h, 5h, 24h, and 48h after cell stimulation). We performed bulk ATAC-seq analysis because, contrary to the single-cell version of this technique, this approach can reliably identify global systemic changes in chromatin structure (Chen et al., 2019). In order to identify relevant DNA regions, we applied a stringent filter based on the reproducible detection of accessibility in all three donors (Aranyi et al., 2016) (see **Table S4** for donor-related information). Performing ATAC-seq on 5000 cells ensured that the detected accessible DNA regions (peaks) are present in a substantial fraction of cells. Indeed, accessible sites present in individual, or a small number of cells, could not be differentiated from the technical noise.

We found a large number of ATAC-seq peaks in cells at 0h (**Figure 4A**). The number of accessible DNA regions further increased by 10-12% between 0h and 5h around the transcription start sites/promoters (TSS), in the introns and exons, but not in the intergenic regions, then decreased gradually at relatively slow rate over the next 48h (**Figure 4B**). The time-dependent decrease in the number of ATAC-seq peaks varied with their genomic location (**Figure 4B**). While the number of peaks in distal intergenic regions was halved between 5h and 48h, the decrease in the other locations was less significant (**Figure 4B**). In particular, the number of peaks in TSS/promoter regions only dropped by 15% between 0h and 48h indicating that these promoters became inaccessible. The significant number of peaks that appear or disappear indicate a rapid global dynamical change of the chromatin structure.

**Figure 4.**
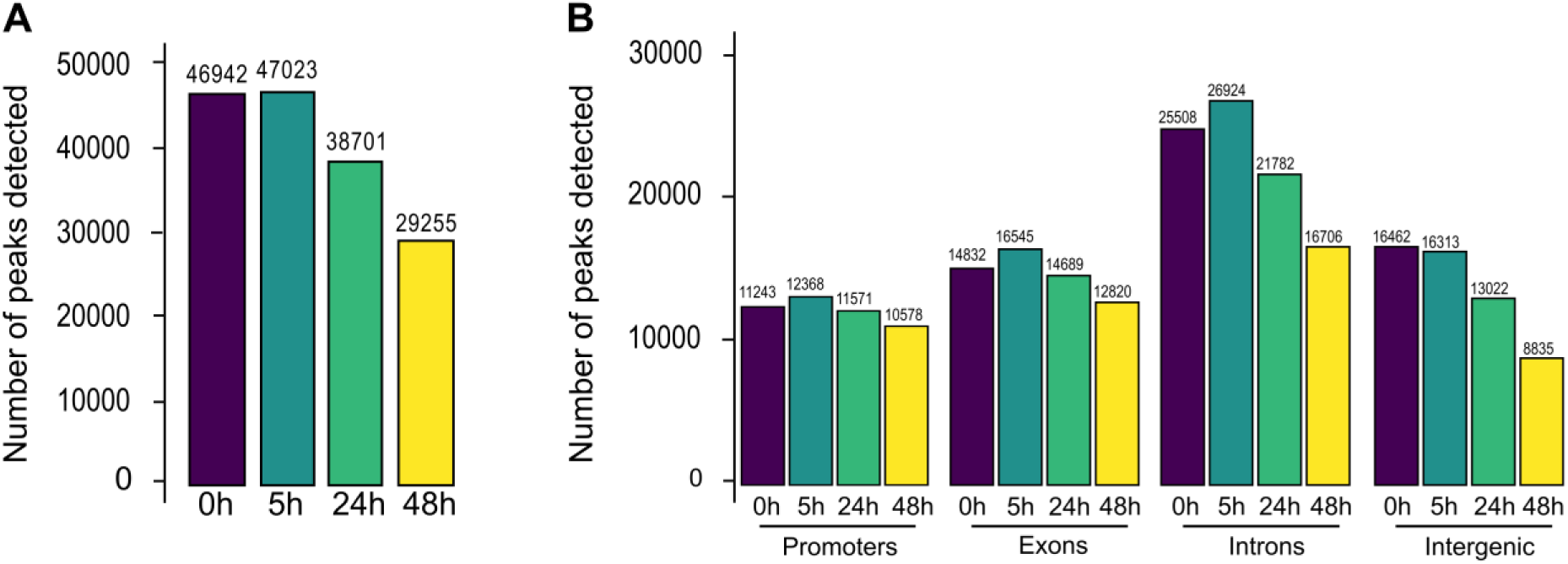
Chromatin dynamics as detected by ATAC-seq. (A) Total number of accessible regions (peaks) at 4 different time points. (B) Number of peaks in different genomic elements. A single peak may count for two categories, except for the intergenic category defined by the exclusion of all the others.

However, the overall tendency emerges from the sum of individual peak dynamics. Therefore, we have analyzed the changes of individual peaks. First, we estimated the changes in the size of the peaks that were present at least at two consecutive time points. As a proxy for the size of a peak, we used the number of sequenced reads that define it. The increase or decrease in read counts for the same ATAC peak between two consecutive time points was used to assess the tendency of the chromatin to open or close, respectively. We calculated the log-fold changes of the number of reads of each peak for time intervals and the associated p-values and represented them as volcano plots (**Figure 5A**). We observed a tendency for the peaks already present at 0h to further increase in accessibility by 5h, in particular peaks located in the TSS regions (blue dots in **Figure 5A**). During this period, accessibility was altered at 17% of the total number of elements detected in our analysis (9,045 out of 53,797). Between 5h and 24h, 15% of the peaks (7,505 out of 50,936) displayed significant change, with approximately equal proportions of increased and decreased subsets. However, between 24h and 48h, only 48 out of 40,248 peaks showed differential read counts, again with roughly equal proportions of increased and decreased peaks (**Figure 5A**). Overall, our ATAC-seq analysis shows that most of the changes in accessibility occurred during the first 24 hours (**Figure 5A**). First, new genomic elements become accessible and others already open become more accessible during the first 5 hours. Then, the trend is reversed: both the number and size of ATAC-seq peaks decreased between 5h and 24h. The latter trend was maintained, albeit at a lesser degree, between 24h and 48h. Although this analysis provides a quantitative assessment of the changes between two time points, it gives no information on the evolution dynamics of individual peaks. Therefore, we plotted the size of each peak at each time. This representation gives a precise account of the changes at each peak. On **Figure 5B**, we represented the peaks detected in promoters and intergenic regions at all four time points. The majority (75%) of the promoter-associated peaks belong to this category. In the intergenic region, only 27% of the peaks are detected at all time points. In both cases, the size of the peaks increased rapidly between 5h and 24h and gradually decreased between 24h and 48h (**Figure 5B**). The peaks that displayed more complex dynamics are represented on **Figure S3**; either they appeared later than 5h or disappeared completely at some stage. However, in both categories, the general tendency to decrease remained the same.

**Figure 5.**
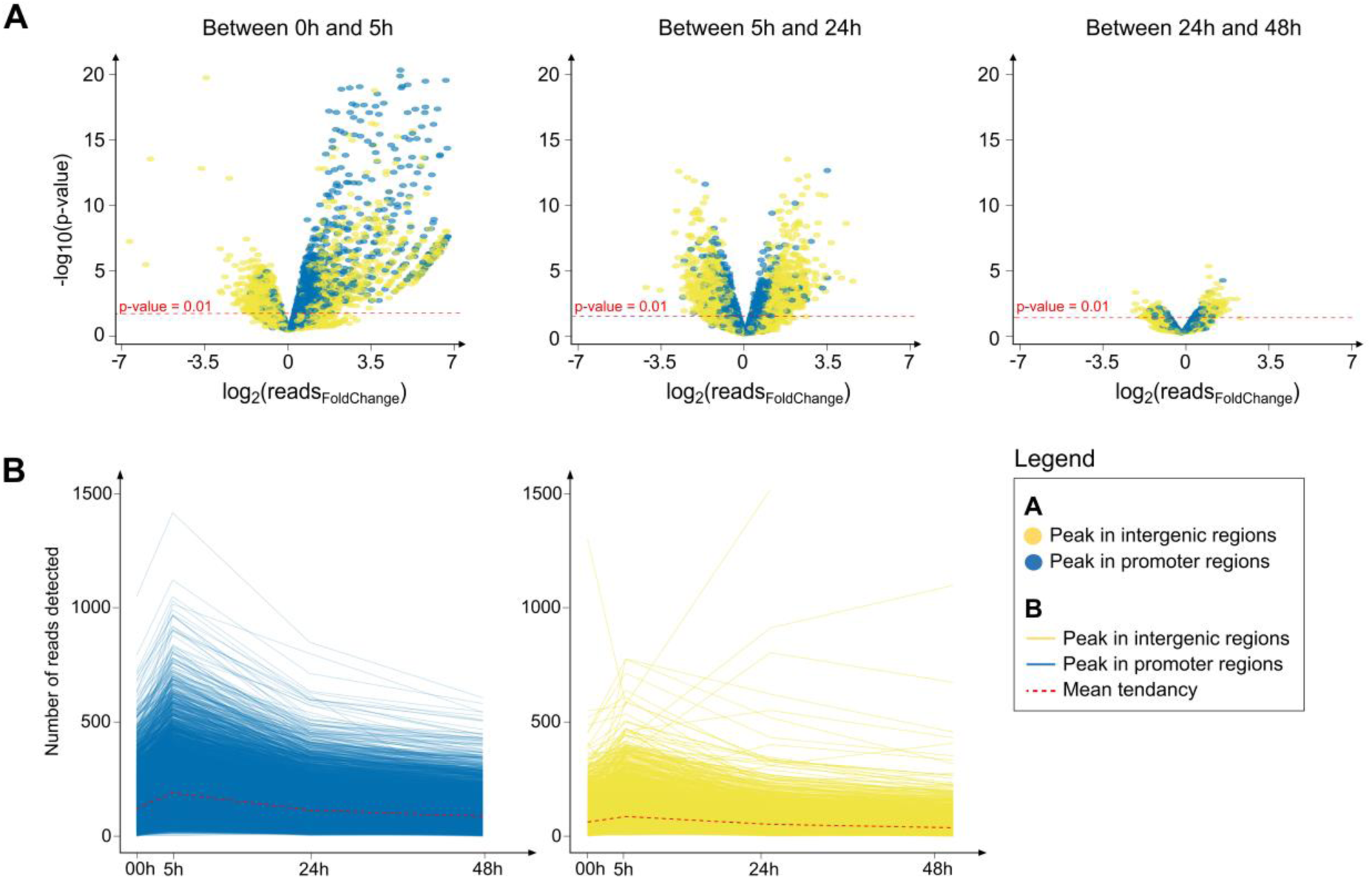
Rapid decompaction and slow re-compaction of the chromatin. (A) Quantitative analysis of the peak sizes detected at two consecutive time points. Peaks in promoter regions are highlighted in blue and in intergenic regions in yellow. Note the significant increase in size (accessibility) between 0h and 5h and the decreasing number of changes after 24h. Details about how the differential accessibility has been calculated are given in STAR Materials and Methods. (B) Evolution of the ATAC peaks in promoters (blue, left panel) and intergenic regions (yellow, right panel). The size of each ATAC peak is plotted for every time point. Each line connects the points corresponding to the ATAC peak detected at the same genomic position. Only the peaks detected at each time point are represented.

To investigate the potential functional importance of the gene promoter accessibility, we analyzed the occurrence of various transcription factors binding site (TFBS) motifs in the accessible DNA regions. We observed that many of the TFBSs of factors known to play a role in hematopoiesis, such as RUNX1, ERG, PU.1 and FLI1, were highly accessible at 0h and remained detectable at a similar level up to 48h (**Figure S4**). We also noted that CTCF (CCCTC-binding factor) binding sites were detected more than five times more frequently in the accessible regions than expected on the basis of their frequency in the genome. Indeed, CTCF is known to play a key role of chromatin remodeling and loop formation in general (Ohlsson et al., 2010), but also more specifically in the hematopoietic lineage (Kieffer-Kwon et al., 2017).

### Chromatin decompaction precedes transcriptional burst

In order to elucidate how the dynamics of chromatin accessibility and the differential gene expressions were related during the critical state transition, we combined the scRNA-seq and the ATAC-seq data (see Materials and Methods). Comparison of scRNA-seq and ATAC-seq analysis in **Figure 1 and 5** shows that the wave of global chromatin opening of gene promoter/transcription start sites (TSSs) precedes the wave of changes in gene transcription. To make sense of this, we first examined how changes in the accessibility gene promoter regions are related to changes in the gene expression. We grouped the promoters in 4 groups: “open-open”, “open-close”, “close-close” and “close-open”, depending on the presence or absence of ATAC-seq peaks at the given promoter at 5h and 24h, respectively (**Figure 6A**). The period between 5h and 24h is particularly interesting and important, because most of the changes in gene expression occur at this stage. We then identified the genes controlled by each promoter using the Regulatory Circuit resource (see Materials and Methods). Finally, we examined the distribution of DE and non-DE genes among the four classes of promoter configuration (i.e. open-open, open-close, close-open, and close-close). Strikingly, 74,2% of DE genes (p < 10^-4^) had a promoter with “open-open” configuration between 5h and 24h (**Figure 6A**), meaning that their promoter was already accessible 5h after cell stimulation, but long before the burst of transcription and they remained so 24h later (**Figure 6A**). This is significantly higher than the proportion of the DE genes in the other categories of promoter configuration as assessed by enrichment analysis. The same classification between 24h and 48h revealed similar repartition of DE genes among categories of promoter configuration (**Figure 6B**). Particularly, more than 60% of DE genes are associated with the “Open-Open” promoter configuration. However, the total number of DE genes is much lower during this period (n = 1849) compared to the first 24 hours (n = 6230) and statistical tests did not reveal any significant overrepresentation of gene categories (**Figure 6B**).

**Figure 6.**
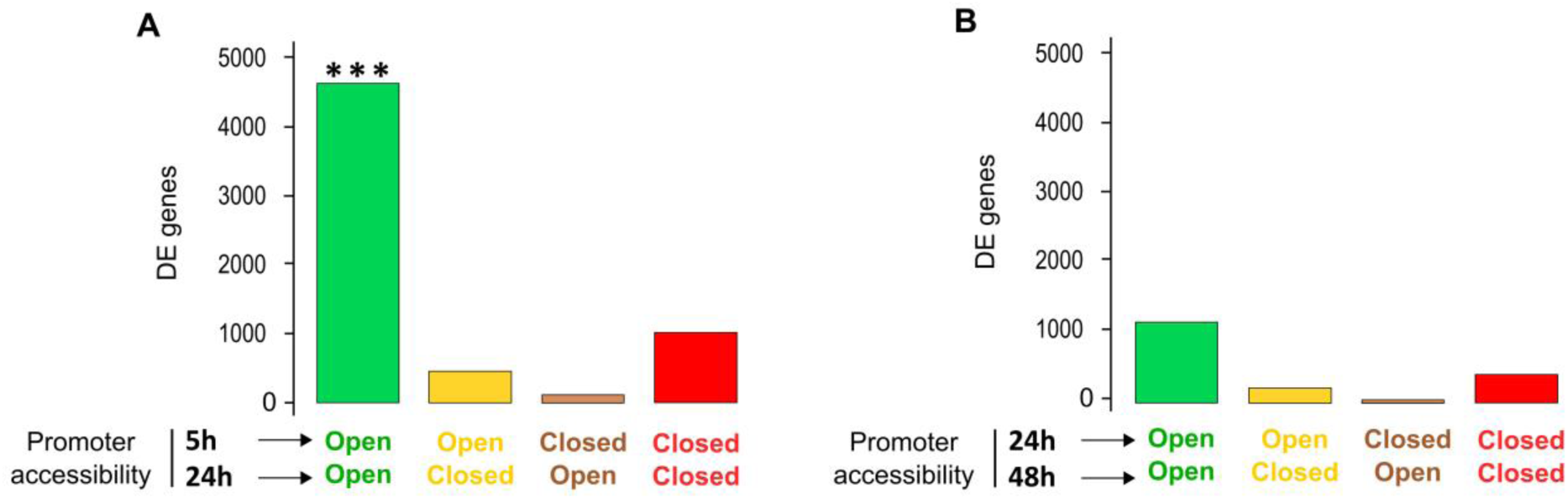
Promoters of differentially expressed genes are continuously accessible. (A) Total number of differentially expressed genes (DE genes) as a function of their promoter accessibility at 5h and 24h. (B) Total number of differentially expressed genes (DE genes) as a function of their promoter accessibility at 24h and 48h. Note that differential genes expression is significantly associated to the Open-Open promoter configuration between 5h and 24h.

We finally examined how alterations of TF expression influenced target gene transcription in combination with the DNA accessibility of the promoter. To do so, we further categorized the DE genes assigned to the four groups according to the chromatin configuration of their promoters “open-open”, “open-close”, “close-close” and “close-open”–depending on whether they were targeted by DE or non-DE TFs, as determined previously (see Single-cell gene expression analysis using RNA-seq). Between 5h and 24h, we found a significantly higher proportion of DE genes in the category DE-TF with “open-open” chromatin configuration than in all other categories (46%; p < 2.5x10^-7^) (**Figure 7A**). In comparison, only 33% of the DE genes were in the non-DE-TF category with “open-open” chromatin. We performed the similar analysis on the ATAC-seq and sc-RNA-seq results obtained at 24h and 48h (**Figure 7B**). No significant enrichment was found for the other categories. The highest fraction of DE genes was found to be associated to the “Open-Open” promoter configuration category with non-DE-TF (51%) (**Figure 7B**).

**Figure 7.**
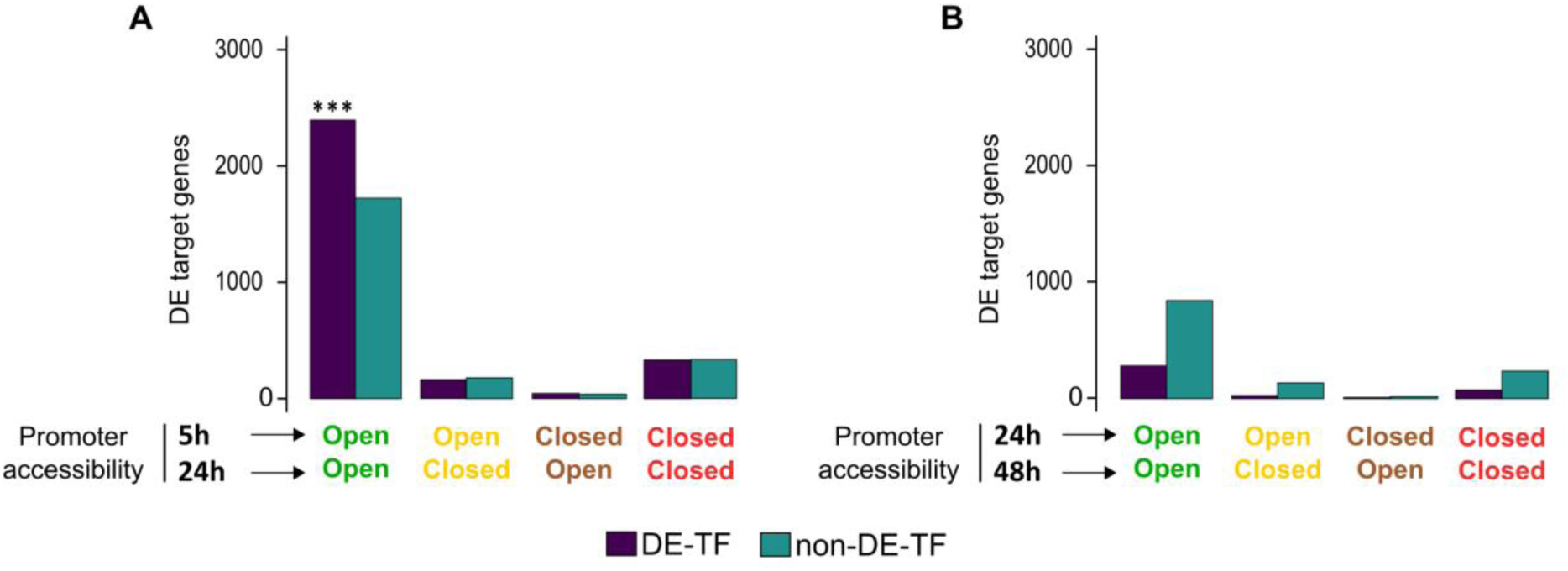
Genes targeted by differentially expressed TF-s are expressed differentially if their promoters are continuously accessible. (A) Total number of differentially expressed genes (DE genes) targeted by differentially expressed (DE-TF) or non-differentially expressed (non-DE-TF) transcription factor-coding genes as a function of their promoter accessibility at 5h and 24h. (B) The same as in A for 24h and 48h.

Taken together, the integration of gene expression and chromatin accessibility data shed light on the chronology of transcriptional regulation in the CD34+ cells. We observed that genome-wide chromatin opening precedes the multilineage-type mixed hyper-expression of a very large number of genes. After 48h, both gene hyper-expression and the number of accessible promoters and extragenic sites started to decrease concomitantly with the emergence of distinct cell populations with particular gene expression patterns.

## Discussion

*In vitro* cultured human cord blood derived CD34+ cells are usually considered as a heterogenous population of cells. Recent studies demonstrated that this heterogeneity is not the result of the mixture of different cell types or subsets, but a population of cells with a wide distribution of gene expression patterns (Velten et al., 2017) that fluctuate between transitory states, generating morphological and transcriptional instability (Moussy et al., 2017). During the first cell cycle, each cell displays a rather distinct gene expression pattern but similar morphology. By 48 to 72 hours, one can observe the emergence of two different cellular morphologies and two different characteristic transcription profiles (Moussy et al., 2017). This observation prompted us to investigate this critical window of time in more details.

Using ATAC-seq, we demonstrated that concomitantly with the cell stimulation the chromatin undergoes very rapid global decompaction followed by gradual condensation. The process of decompaction reached a maximum as early as 5h after the stimulation of the cells and made most of the gene promoters in the genome accessible. The opposite process of closure is slow and gradual.

Importantly, the rise-and-fall in chromatin opening precede and overlap with a rise-and-fall in transcriptional activity peaking at 24-48h. Indeed, the variety in transcribed genes and the number of mRNA molecules per gene was the lowest at 5h – the first time point tested for scRNA-seq – but both increased sharply at 24h, reached a plateau between 48h and 72h and decreased at 96h (**Figure 1B and 1C**). The 5h-to-48h period corresponds to the multilineage-primed stage of the CD34+ cells that precedes the emergence of the first signs of characteristic gene expression patterns accompanying differentiation (Moussy et al., 2017).

Progress through a transitional cell state marked by the rise-and-fall in transcriptional uncertainty and a concomitant rise-and-fall of cell-to-cell variability was previously reported as a universal feature of cells during the initial phases of the fate commitment process (Gao et al., 2020). We show here using CD34+ cells that the global increase in transcription most likely arises as a consequence of a widespread and non-specific chromatin opening that makes widely accessible more than 50% of gene promoters in the genome. Importantly, the number of gene promoters becoming accessible largely exceeds the number of genes that are actually transcribed in each cell (**Figure 1B and 4B**), pointing to a strong stochastic component in the establishment of the multilineage primed expression state. Coherent transcription profiles emerge from this heterogeneous transitory state concomitantly with the gradual chromatin compaction. A significant fraction of gene promoters (16%) and intergenic sites (46%) in the genome become inaccessible again between 5h and 48h (**Figure 4B**). The stabilization of the transcriptome is presumably the consequence of these chromatin changes. Some promoters gradually become repressed by chromatin closing, while others are stabilized in an open chromatin configuration. The role of TFs appears crucial at this stage. Indeed, between 5h and 24h the increase of the transcription of TF-encoding genes correlated with the similar increase of their target genes with accessible promoters. Changes of the expression of the TF-encoding genes do not alter the target gene expression if their promoters are in “closed” chromatin configuration around the TSS (**Figure 7A and 7B**), indicating that chromatin accessibility plays a permissive or gating role for TF action. Since the number of the open promoters is higher at the beginning of the process than the number of expressed genes, a competition for the available TFs among accessible promoters may explain the transcriptional and phenotypic fluctuations observed during this period (Moussy et al., 2017). These fluctuations cease when the transcriptome is stabilized (Moussy et al., 2017). The role of TFs may be crucial during the second phase, because their binding may keep the target genes transcribed and prevent the closing of the chromatin. The proposed scenario of general non-specific chromatin destabilization followed by a selective repression of the genes is also supported by the observations showing that the inhibition of chromatin compaction using valproic acid (VPA), a histone deacetylase inhibitor, can maintain the multilineage-primed state with promiscuous transcription profile for a long period (Chaurasia et al., 2014; Moussy et al., 2019, 2017). The removal of VPA allows defined transcriptome profiles to be established (Moussy et al., 2019). Therefore, chromatin structural changes appear to be causally involved both in the generation of a non-specific multilineage-primed transcriptional state and the stabilization of the cell fate choice.

Recent mechanistic studies in various cellular systems support our model. For example, a recent study of human fetal hematopoietic cells demonstrated that extensive epigenetic, but not transcriptional priming of HSC/MPPs, occurs prior to lineage commitment (Ranzoni et al., 2020). In another study, monitoring the alterations in the chromatin structure and the nuclear architecture during B cell activation revealed that as quiescent lymphocytes encounter antigens, they rapidly decondense chromatin by spreading nucleosomes from the nuclear matrix to the entire nucleoplasm, decompacting chromatin clusters into mononucleosome fibers, and strengthening their nuclear architecture by creating new CTCF loops and contact domains. The global decompaction and loop formation require Myc, constant energy input, histone acetylation, and is accompanied by an increase in regulatory DNA interactions and gene expression (Kieffer-Kwon et al., 2017). Studies on hair bulb stem cells also showed that changes in chromatin accessibility precede gene expression changes and lineage commitment (Ma et al., 2020). Similarly, the loss of DNA methylation has been shown to be essential for the establishment of chromatin accessibility that determines differential transcription factor binding in neural stem and progenitor cells. Following the differentiation into glial cells, new methylation is acquired to maintain the identity of glial cells by silencing neuronal genes (Sanosaka et al., 2017). Furthermore, in human cells, most changes during differentiation arise from dramatic redistributions of repressive H3K9me3 and H3K27me3 marks, which form blocks that significantly expand in differentiated cells (Hawkins et al., 2010).

While the rapid and non-specific opening of the chromatin as a general response to stimulation appears now to be well documented, it is of particular importance for further understanding to investigate the process of transcriptome stabilization and the feedback mechanisms that must accompany the emergence of specific gene expression patterns. In this respect, it may be relevant that a dynamic positive feedback loop between permissive chromatin and translational output has been previously reported for embryonic stem- and in CD34+ cells (Bulut-Karslioglu et al., 2018). It is noteworthy that many of the genes with the most variable expression that contribute significantly to the specification of the emerging transcription patterns are ribosomal protein (RP) coding genes (**Table S2**), thus impacting the process of translation (Guo, 2018). A high degree of RP expression heterogeneity has already been observed in hematopoietic cells, where a small subset of RPs can discriminate cell types belonging to different hematopoietic lineages (Guimaraes and Zavolan, 2016). Therefore, it is possible that, in addition to the TF and promoter interactions, a feedback action of the translational output may also contribute to the stabilization of the chromatin. Analogous feedback regulation has been described in ES cells where the translational output directly promotes a permissive chromatin environment, in part by maintaining the levels of unstable euchromatin (Bulut-Karslioglu et al., 2018). Clearly, the selective stabilization of the chromatin is impacted by many more mechanisms, but their respective roles remain to be clarified.

The observed non-specific chromatin opening and the rise of an equally non-specific gene expression as a first step, followed by a slow relaxation toward a defined gene expression pattern and chromatin stabilization, brings a new perspective to our understanding of how cell fate commitment is initiated. According to the conventional view, a switch-like activation of fate-specifying genes, followed by a cascade of activation of specific downstream targets determines cell fate. This view is not compatible with the observations reported here. We propose an alternative model where stochastic and highly variable expression profile of multilineage-primed transitory stage emerges as a rapid but non-specific response to a substantial change in the cell’s environment. This reaction is analogous to the physiological stress response whose role is to prepare the organism to meet new and unforeseen circumstances (Braun, 2015). In the case of the cells, we observe a general and non-specific opening of the chromatin that lifts the transcription repression and permits targeted interactions between TFs and gene promoters and enhancers. The quasi-random activation of genes in a cell under stressful conditions generates a potential of a variety of phenotypic traits in the cell. Some of these traits promote the cell’s survival under selective pressures imposed by the evolving microenvironment, and they are gradually and selectively stabilized by feedback mechanisms. All these mechanisms are not yet identified, but explicit and testable hypotheses have been made on their nature (Paldi, 2003; Páldi, 2020).

Overall, fate commitment of the CD34+ cells can be viewed as a continuous iterative process of constrained optimization of the cell phenotype, a kind of “learning process” that is accomplished by the cell through interactions and cooperation with the surrounding cells and environment. This way to conceptualize the question of fate commitment has been theorized long ago (Kupiec, 1997, 1996; Paldi, 2012), and now it is supported by an increasing number single-cell experimental studies (Gao et al., 2020; Hu et al., 1997; Mojtahedi et al., 2016; Moussy et al., 2017; Richard et al., 2016).

## Acknowledgements

The authors are grateful to Olivier Gandrillon, Camille Fourneaux and François Delhommeau for helpful discussions and Sunil Laxman, Takuya Imamura and Olivier Gandrillon for the critical reading of the manuscript. The authors are also grateful to Sophie Foulon for her help in scRNA sequencing protocol design.

This work was supported by EPHE (11REC/BIMO), ANR grant ANR-17CE12-0031-01 «SinCity ».

## Author contribution

AP, AM, RP and JFD designed the study.

RP, AM, LR and SC conducted the experiments.

AM, RS, RG and NPG performed CALISTA analysis.

RP, DS, RS, SC and RG analyzed the ATAC-seq data.

RP, AM, LR, SC, RS, RG, NPG, DS, GC, JC, GF and AP analyzed the results and performed statistical analysis.

RP, AM, GF and LR prepared the figures.

AP, RP, GF and LR wrote the paper with the help of their colleagues.

## Declaration of interests

The authors declare no competing interests.

## Materials and Methods

### Cell culture

Umbilical cord blood from anonymous healthy donors was obtained from Centre Hospitalier Sud Francilien, Evry, France or from AP-HP, Hôpital Saint-Louis, Unité de Thérapie Cellulaire, CRB-Banque de Sang de Cordon, Paris, France (Authorization number: AC-2016-2759). Mononuclear cells were isolated from cord blood fractions by density centrifugation using Ficoll (Biocoll, Merck Millipore, Burlington, Massachusetts). Human CD34+ cells were then enriched in the sample by immunomagnetic beads using an AutoMACSpro (Miltenyi Biotec, Bergisch Gladbach, Germany). After collection, enriched CD34+ cells were frozen in a cryopreservation medium containing 90% of fetal bovine serum (Eurobio, Les Ulis, France) and 10% of dimethylsulfoxide (Sigma, Saint-Louis, Missouri) and stored in liquid nitrogen.

After thawing, the CD34+ cells were cultured in a 96-well plate in a humidified 5% CO2 incubator at 37°C. Cells were cultured in prestimulation medium made of X-Vivo (Lonza, Basel, Switzerland) supplemented with penicillin/streptomycin (respectively 100U/mL and 100ug/mL - Gibco, Thermofisher Scientific, Waltham, Massachussetts), 50 ng/ml h-FLT3, 25 ng/ml h-SCF, 25 ng/ml h-TPO, 10 ng/ml h-IL3 (Miltenyi Biotec, Bergisch Gladbach, Germany) final concentration.

### Fast-ATAC-seq

We used Fast ATAC-seq with minor modifications. This protocol was optimized for blood cells (Corces et al., 2016). Prior to transposition, cells were marked with 7AAD, and dead cells were removed by FACS (Beckman Coulter, Brea, California). Removing dead cells is an important parameter to ensure clear nucleosome patterns and to improve signal to noise ratio. 5000 living cells were used at each time point. A one-step gentle membrane permeabilization and DNA transposition was performed by adding 50ul transposition mixture (25 uL TD buffer 2X, 2,5uL of transposase TDE1 (Illumina, San Diego, California), 0,5 uL digitonin 0,1% (Promega, Madison, Wisconsin) and 22 uL water) to the cell pellets and by incubating at 37°C for 30 minutes under agitation. Obtained Transposed DNA were then purified using MinElute PCR Purification Kit (Qiagen, Hilden, Germany) and preamplified using Nextera barcoded primers (Illumina, San Diego, California) and NEBNext High-Fidelity 2xPCR Master Mix (New England Biolabs, Ipswich, Massachusetts) for 5 cycles. A quantitative PCR amplification was made on 5uL of the sample with SYBR Green to determine the number of additional cycles in order to generate libraries with a minimal number of PCR cycles and to limit PCR bias (according (Corces et al., 2016)). Appropriate number of PCR cycles were applied on the rest of the pre-amplified samples. PCR fragments were purified with MinElute PCR Purification Kit (Qiagen, Hilden, Germany) to get rid of unused primers. A supplemental purification step was performed using Ampure beads kit (Beckman Coulter, Brea, California) to size-select DNA fragments ranging between 100 and 700 pb. ATAC-seq libraries were checked for quality using Bioanalyzer (Agilent, Santa Clara, California) prior to sequencing and sequenced in paired-end mode (2x50bp) on the Illumina HiSeq2500 platform.

### Single-cell RNA sequencing adapted from MARS-seq

To perform scRNA-seq, we adapted the MARS-seq protocol (Massively parallel single-cell RNA sequencing) (Jaitin et al., 2014). CD34+ cells were stained with 7AAD to only work living cells and cells were isolated by FACS. Individual cells were sorted into 96-well plates containing 4uL of lysis buffer with specific barcoded RT primers (final concentration: 0,2% Triton, 0,4 U/μL RNaseOUT (Thermofisher Scientific, Waltham, Massachussetts), 400nM idx_RT_primers). Idx_RT_primers contain a T7 RNA polymerase promoter for further *in vitro* transcription (IVT), single cell barcodes for subsequent de-multiplexing and unique molecular identifiers (UMIs) allowing correction for amplification biases (**Table S6**). After cell sorting, plates were immediately centrifuged and put into dry ice before storage at -80°C preceding the reverse transcription (RT). To open RNA secondary structure, plates containing single cells were incubated at 72°C for 3 minutes and immediately put in ice. 4μL of RT mix were added in each well (final concentration of RT mix: 20mM DTT, 2mM dNTP, 2X First stranded buffer, 5 U/μL Superscript III RT enzyme, 10% (W/V) PEG 8000). PEG8000 was added in the RT mix because it has been shown that it can increase the cDNA yield in scRNA sequencing (Bagnoli et al., 2018). ERCC RNA spike-in mix (Thermofisher Scientific, Waltham, Massachussetts) was also added to the solution for further amplification quality filtering (dilution 1/40.10e7). The plate was then put into thermocycler (thermocycler program: 42°C-2min, 50°C-50min, 85°C-5min, 4°C hold).

After first retro-transcription, samples were pooled (see (Jaitin et al., 2014)) and ExonucleaseI digestion was performed, followed by 1,2X AMpure beads purification kit (Beckman Coulter, Brea, California) to keep only retro-transcribed single strand cDNA. Samples were eluted in 17μL of 10mM Tris-HCl, pH=7,5. Second strand cDNA synthesis (SSS) using NEBNext mRNA second strand synthesis module kit was then performed (SSS mix: 2μL 10x SSS buffer, 1μL SSS enzyme; thermocycler program: 16°C-150min, 65°C-20min, 4°C hold). Obtained cDNA was linearly amplified by overnight IVT (HighScribe T7 High Yield RNA synthesis, NEB) at 37°C under T7 promoter. The product was purified with 1,3X Ampure beads and eluted in 10μL of 10mM Tris-HCl, 0,1mM EDTA. 9μL of amplified RNA were then enzymatically fragmented with 1uL of 10x RNA fragmentation reagents (Thermofisher Scientific, Waltham, Massachussetts) in 70°C for 3 min. The fragmentation was stopped with 34μL of STOP mix (1,2uL Stop solution, 26,4μL AMpure beads, 9,8uL TE) and samples were purified. Differing from original MARSseq protocol, the second RT was done with primers (P5N6_XXXX) containing random hexamers and specific barcode (**Table S6**) to distinguish the different plates (*ie.* times) (final concentration: 5mM DTT, 500uM dNTP, 10uM P5N6_XXXX, 1X First stranded buffer, 10U/μL Superscript III RT enzyme, 2U/μL RNaseOUT; thermocycler program: 25°C 5min, 55°C 20min, 70°C 15min, 4°C hold). cDNA was purified with 1,2x AMpure beads and eluted in 10μL.

As for ATAC-seq, the appropriate number of PCR cycles was determined using a fraction of the library with SYBR Green based qPCR as described in (Zilionis et al., 2017) (final concentration: 1x Kapa Hifi HotSTart PCR mix, 1x SybrGreen, 0,5μM mix primer P5.Rd1/P7.Rd2; Thermocycler program: 95°C 3min – 40cycles: 98°C 20sec, 57°C 30sec, 72°C 40sec – 72°C 5min, 4°C hold). After PCR amplification, libraries were purified with 0,7x AMpure beads. Libraries were checked for quality, using Bioanalyzer HighSensitivity DNA (Agilent, Santa Clara, California) prior to sequencing. Libraries were finally sequenced in paired-end mode (2x50bp) on Illumina HiSeq2500 platform.

Idx RT primers: TTTTTTTTTTTTTTTTTTTTN = poly-T allowing matching with mRNA poly-A tail, NNNN = 4 bases UMI (randomly generated), XXXXXX = 6 bases cell barcode (**Table S6**). The rest of the sequence consists of a PCR adaptor and a T7 promoter sequence for further IVT amplification. P5N6 XXX: NNNNNN = random hexamer allowing the capture of the fragmented IVT amplified RNA, XXXX = 4 bases “plate barcode” (**Table S6**). The rest of the sequence consists of a PCR adaptor. P5.Rd1/P7.Rd2 : P5 and P7 Illumina sequencing adaptors.

### Bioinformatic analysis

#### Single-cell RNA-seq (scRNA-seq)

##### Raw data processing

Cell and plate barcode demultiplexing steps were accomplished under strict selection criteria with the following command:

*< cutadapt -q 30 -e 0 -m 30:20 --no-trim --no-indels --pair-filter = any >*

Sequence for both barcodes (cells and time) sequences are given in **Table S6**.

ERCC mapping was performed using bowtie2 (Langmead and Salzberg, 2012) on ERCC known sequences and regular mapping was performed using STAR (Dobin et al., 2013) on the reference genome version hg19 and aligned reads annotated. After quality filtering, reads and UMIs count per gene and ERCC were calculated for expression analysis.

##### Cell and gene filtering

Chromosome Y was removed from the analysis to avoid unwanted effects and only protein coding genes were kept for further analysis. Cells with less than 80 000 total reads were removed, as well as cells with more than 10% of reads corresponding to mitochondrial RNA. To reduce undesired effect due to PCR non-linear amplification, ERCC spikes were used to assess the linearity of amplification. Pearson correlation coefficient was calculated for each cell, and only cells above 0,6 were retained. For each cell remaining, genes were defined as detectable if at least two cells contained more than a single UMI (=transcript) and a minimum of 5 reads in total.

##### Single-cell clustering and variability analysis

Clustering analysis was performed with CALISTA (Clustering and Lineage Inference in Single-Cell Transcriptional Analysis) (Papili Gao et al., 2020), a numerically efficient and highly scalable toolbox for end-to-end analysis of single-cell transcriptomic profiles. This approach includes single-cell mRNA counts in a probabilistic distribution function associated with stochastic gene transcriptional bursts and random technical dropout events. In the data pre-processing, we removed cells with more than 95% of zero expression values and then selected the top 200 most informative genes for further analysis. The optimal number of clusters was chosen to be five based on the eigengap plot (see (Papili Gao et al., 2020) for more details).

##### WGCNA

We applied Weighted Correlation Network Analysis (WGCNA) (Langfelder and Horvath, 2008) to the mRNA expression data from each donor separately, to identify modules of genes with similar gene transcriptional dynamics. We excluded genes without any detectable expression in all samples. In implementing WGCNA, we set the soft-thresholding power for a scale-free topology index of 0,9. For each module, we calculated the mean expression of genes by averaging the UMI counts from the two donors separately.

##### Enrichment Analysis

We obtained a curated collection of TFs to CAGE-defined promoters to gene isoform mapping for a total of 662 human TFs from the Regulatory Circuits resource (Marbach et al., 2016; Noguchi et al., 2017). In our analysis, we used only TF – Promoter pairs with moderate confidence scores > 0.5. We grouped genes based on whether the relevant TFs demonstrated differential expressions. More specifically, a classification of “changes in TF” was given to any gene in which at least one of its TFs showed a differential expression. Otherwise, a classification of “no change in TF” was assigned. A two-sided Fisher exact test was used to perform over- and under-representation analysis (Agresti, 2007).

#### Bulk ATAC-seq

##### Raw data processing

Tn5 adapters sequences were first trimmed with the following command:

*< cutadapt -q 20 -g “AGATGTGTATAAGAGACAG; max_error_rate=0.1; min_overlap = 10” -A “AGATGTGTATAAGAGACAG; max_error_rate = 0.1; min_overlap = 10” --minimum-length 18 -- times 2 --pair-filter = both >*

Genome alignment (hg19) was performed using Bowtie2 with the following parameters:

*< bowtie2 -x hg19 --no-unal -X 800 >*

Only Paired-End fragments were kept, considering mapping quality (phred score = 30). Duplicated reads were removed using Picard MarkDuplicates tool. In attempt to not bias the signal recovered after peak calling due to multiple donors, all paired-end files were randomly downsampled to 16M reads (without disrupting pairs of reads) as regard to the smallest number of reads detected in the cohort (Donor 1 – 0h, see **Table S4**).

ATAC-seq peaks were then called on those downsampled files using:

*< macs2 callpeak -f BAMPE -g hs -B --broad --broad-cutoff 0.1 --keep-dup all >*

In order to retain only significant accessibility peaks across samples, each list of peaks used in advanced analysis has been defined as the intersection between peaks of the 3 donors tested at the same time point.

##### Peak annotation

Peaks were assigned to genomic regions thanks to a home-made script based on the FindOverlap function from the R package “GenomicRanges” (Lawrence et al., 2013). Genomic elements positions (exons, introns, CpG islands and CTCF) were retrieved from UCSC database (hg19). As for the RNA-seq analysis, promoters regions were retrieved from the online database FANTOM5 (Noguchi et al., 2017). Intergenic category was defined as the exclusion of all other defined categories. No priority has been set across the different genomic elements. Therefore, peaks overlapping several genomic features are counted multiple times, resulting in a total number of peaks across elements exceeding the total number of peaks detected at each time point.

##### Peak differential analysis

DEseq2 tool was used to calculate difference in read count between peaks in two consecutive time points (Love et al., 2014). More precisely, the region considered is defined as the interval formed by the union of two overlapping peaks at *t2* and *t1*.

##### Motif enrichment

Peak motif enrichment analysis was conducted with the tool “findMotifsGenome.pl” from the HOMER software tool suite (Heinz et al., 2010). Background file was generated using an auto-generated list of random regions across the genome (hg19). Motifs were scanned using the total length of our peaks by providing the option *<size given>*.

#### ATAC-seq and scRNA-seq combined analysis (accessibility – expression)

##### Identification of Promoters that have configurational changes

In an effort to identify promoter regions that are affected (and not affected) by configurational changes of the chromatin, we employed the R Bioconductor package “GenomicRanges” (Lawrence et al., 2013). By comparing the peaks overlapping the promoters between two time points (0h – 5h, 5h – 24 h and 24h – 48h), we grouped promoters into 4 possible chromatin accessibility configurations: “open-open”, “open-close”, “close-open”, and “close-close”. We then used the CAGE-defined promoters to gene isoform mapping from the Regulatory Circuits resource (Marbach et al., 2016; Noguchi et al., 2017) to identify promoters that overlap with the peaks of ATAC-seq and their corresponding target genes.

##### Differential gene expression of single-cell RNA sequencing

We computed Z-scores for every gene in each of the two donors between two different time points using the mean and standard deviation of the UMI counts of approximately 100 single cells.

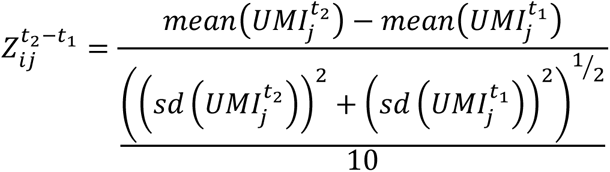

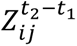 denotes the Z-score of the expression change of gene *j* in donor *i* between time *t*_2_ and *t*_1_. An average Z-score between the two donors was computed and used to identify the set of differentially expressed genes. We selected Z-score thresholds of 2 and –2 (i.e., two standard deviations of change) to designate upregulated and downregulated genes, respectively. Collectively, they represent the set of differentially expressed genes (DE genes).

##### Enrichment Analysis of Combined ATAC-seq and scRNA-seq

For the combined ATAC- and scRNA-seq analysis, we grouped genes into 8 possible groups based on the chromatin accessibility configurations (*i.e.*, one of the following four configurations: “open-open”, “open-close”, “close-open”, and “close-close”) and whether at least one of their TFs coding genes showed differential expression (*i.e.*, one of the following two groups: “DE-TF” and “non-DE-TF”) (**Figure 5C**). As with the analysis of scRNA-seq data, a gene was assigned to the group “DE -TF” when at least one of its TFs showed differential expression; otherwise, the gene was classified as “non-DE-TF”. Note that different isoforms of the same gene can have distinct TSSs that are under the control of different promoters. Thus, a gene might be counted in more than one category in the chromatin accessibility configurations. Consequently, the total sum of the genes in the 8 groups as described above might exceed the total number of genes. A two-sided Fisher exact test was used to perform over- and under-representation analysis (Agresti, 2007).

## Supplemental information

**Figure S1.**
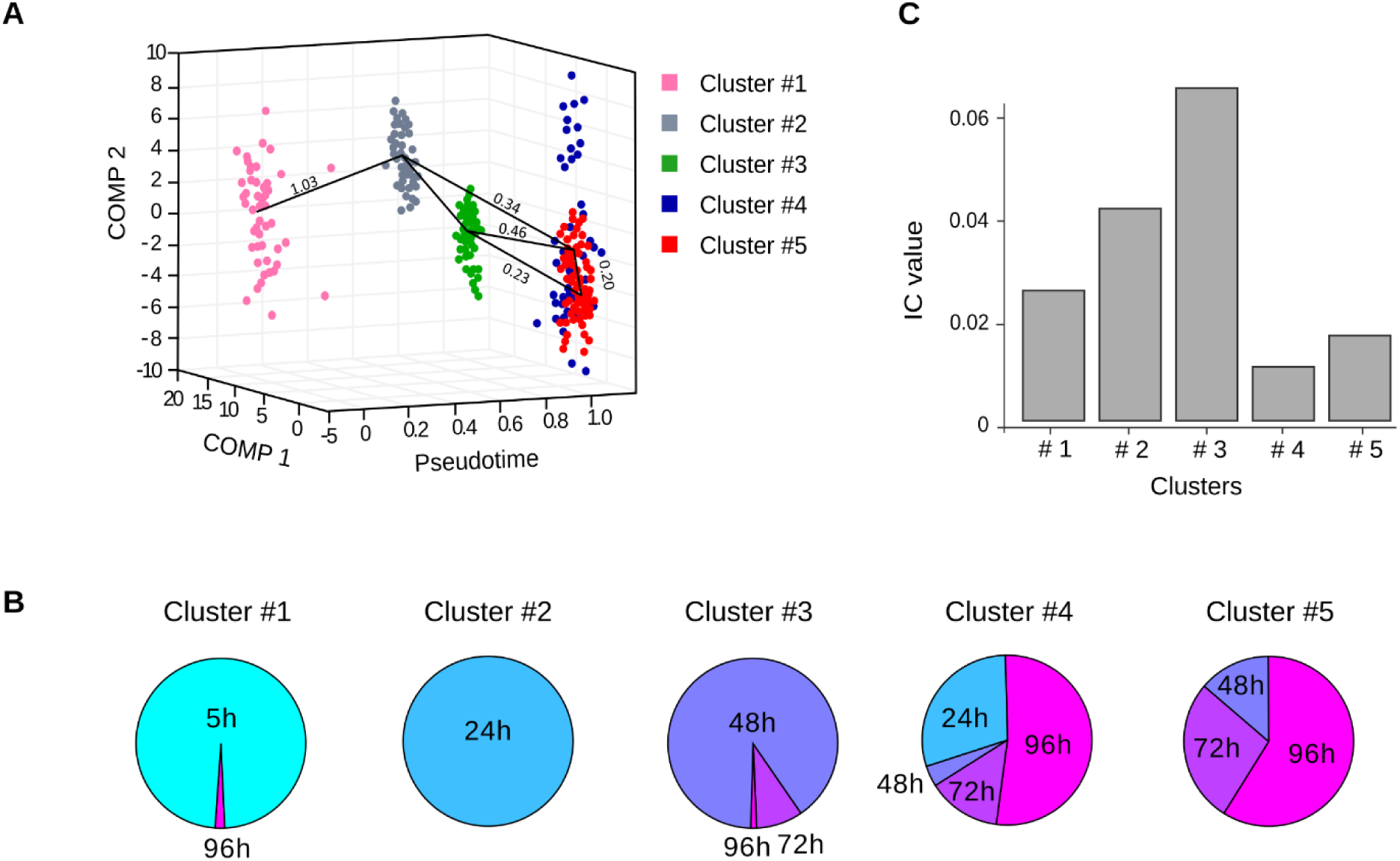
Evolution of transcriptome profiles after cell stimulation in Donor 1. (A) Transcriptome clusters identified by CALISTA for donor 1. Each dot corresponds to a cell in the single-cell transcriptomic dataset sampled at 5h, 24h, 48h, 72h and 96h. The x axis corresponds to the pseudotime, the y-z axes to the first and second principal component (PC). The color code is given in the upper right inset. The transition edges are represented by black plain lines between the clusters and the numbers are “cluster distances”, a likelihood-based measure of dissimilarity (distance) between cell clusters. Note that there are several ways a cell can reach the clusters 3 to 5. The results for Donor 2 are shown on Figure 2. (B) Contribution of the cells collected at different time points to the clusters identified by CALISTA. The mixed composition of the clusters #3 to #5 may reflect the different rates of cell transformation and the multiplicity of cell trajectories. (C) Ic index calculated for each cluster of donor 1 as described in (Mojtahedi et al., 2016). The maximum is reached for cluster #3, indicating a phase of critical transition at 48h. As soon as 24h, but mainly after 48h, cells from cluster #4 and #5 undergo stabilization, leading to a decreasing Ic index value.

**Figure S2.**
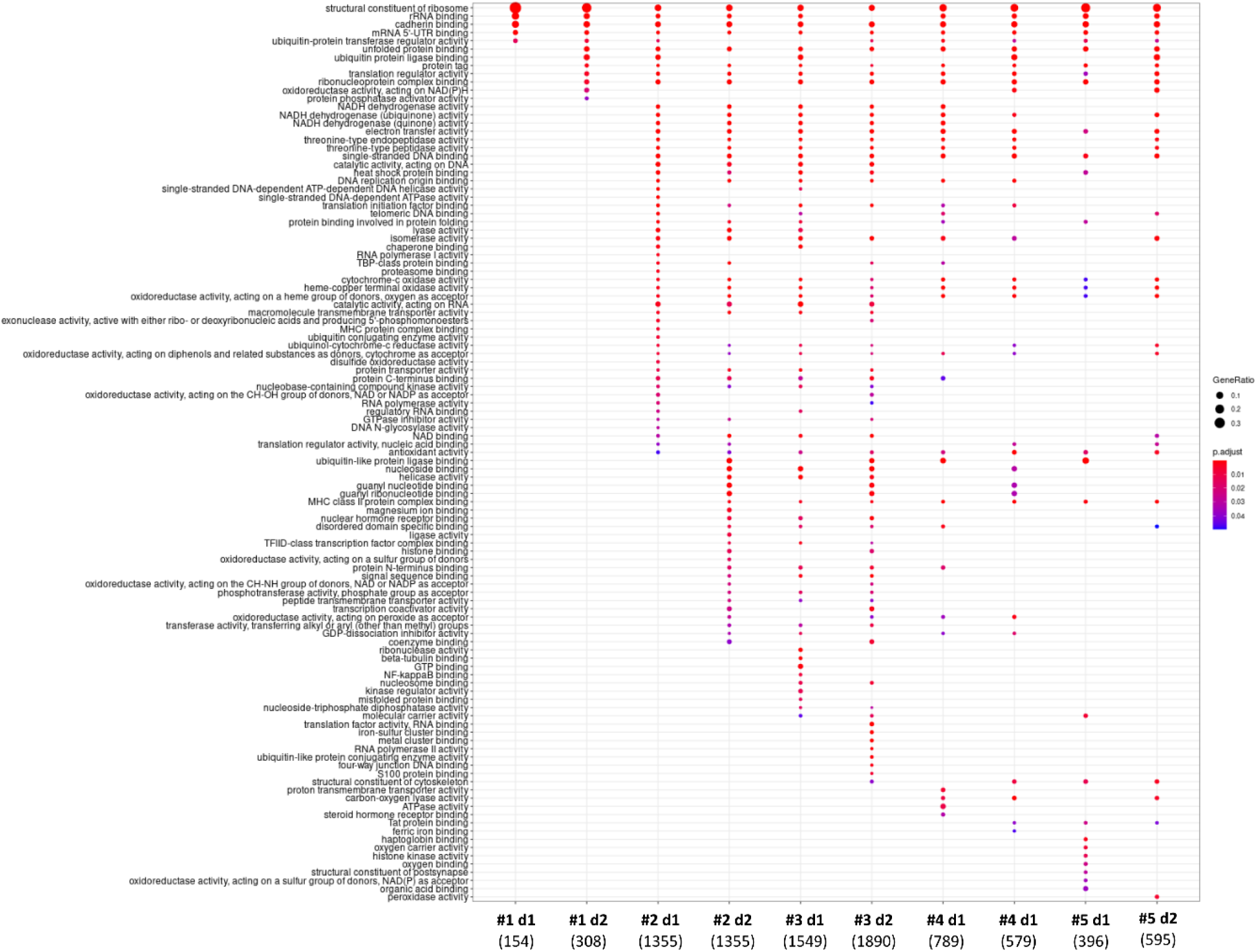
Comparative GO enrichment analysis of clusters in both donors. Top gene ontology categories (GO) found in the clusters of cells determined with CALISTA (p- adj < 0.05). Genes with pairwise gene-gene correlation scores greater than 0.70 were used for the GO analysis. Columns correspond to individual clusters (#) from donor 1 (d1) and 2 (d2). Numbers of genes associated to each cluster are indicated between parentheses under each cluster, on the x-axis. For GO terms associated statistics and entrez gene IDs, see **Table S3**.

**Figure S3.**
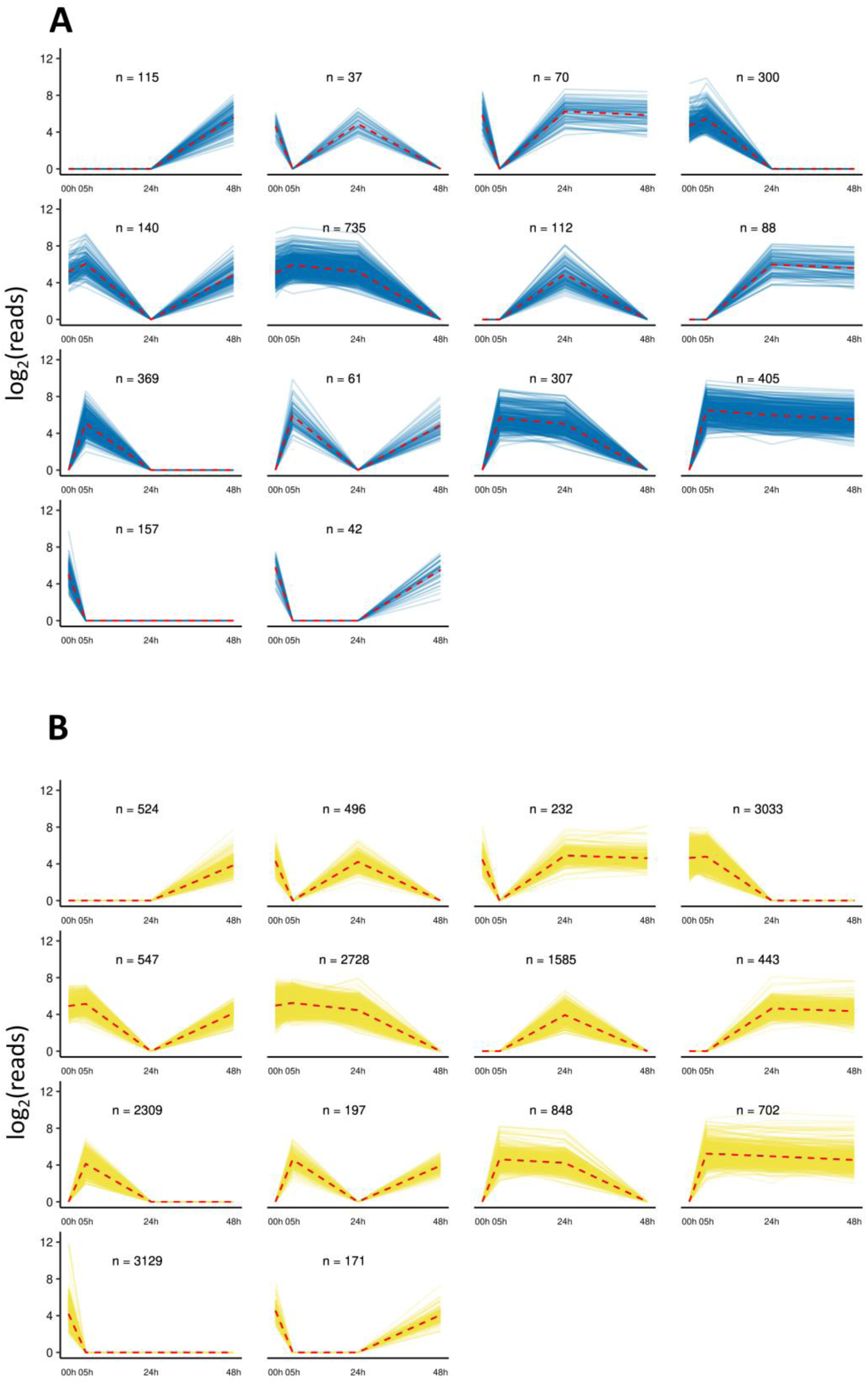
Promoter-associated and intergenic ATAC peaks with complex dynamics. (A) Promoters (blue). (B) intergenic regions (yellow). The size of each ATAC peak is plotted for every time point. Each line connects the points corresponding to the same genomic position where a peak was found.

**Figure S4.**
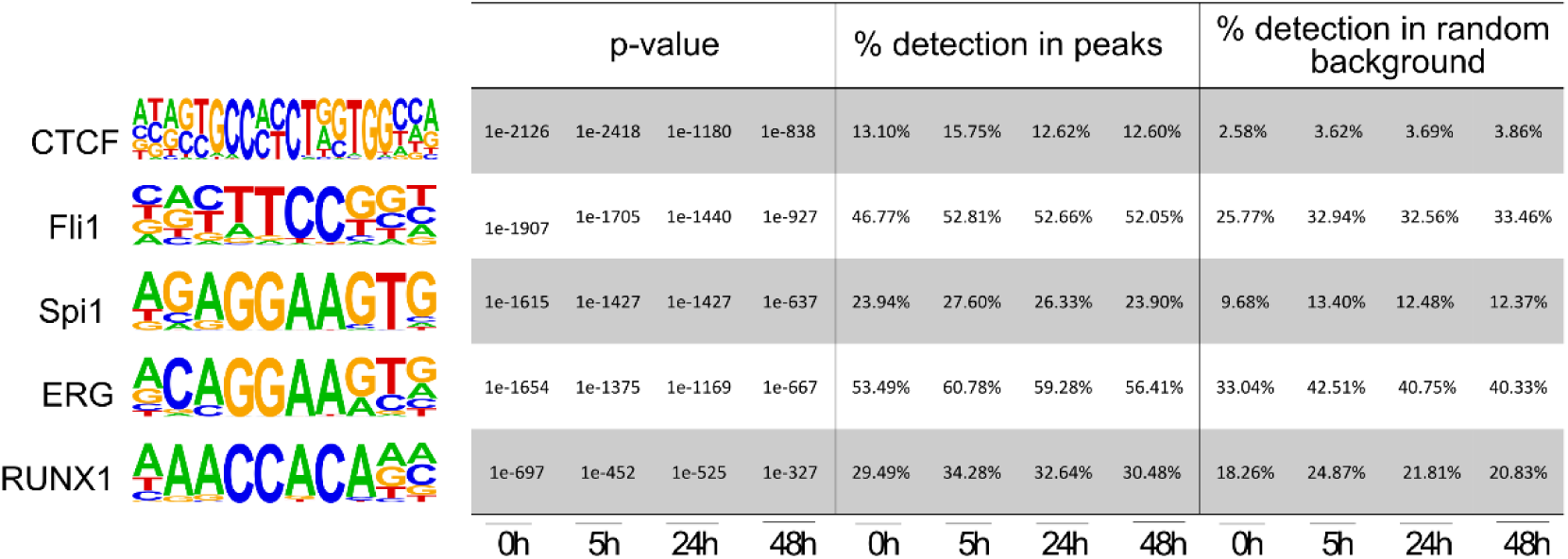
Enrichment of selected known hematopoiesis related transcription factor binding motifs. At each time point, peak sequences were scanned by HOMER for “known motifs”. The motifs five of the factors selected here showed the most significant enrichment. They are well known to be associated with hematopoiesis and chromatin remodeling. For an extensive list of tested motifs and statistics, see **Table S4**.

## Notes

### Competing Interest Statement

The authors have declared no competing interest.

